# A simple optical flow model explains why certain object viewpoints are special

**DOI:** 10.1101/2023.10.02.560466

**Authors:** Emma E.M. Stewart, Roland W. Fleming, Alexander C. Schütz

## Abstract

A core challenge in perception is recognizing objects across the highly variable retinal input that occurs when objects are viewed from different directions (e.g., front *vs* side views). It has long been known that certain views are of particular importance, but it remains unclear why. We reasoned that characterising the computations underlying visual comparisons between objects could explain the privileged status of certain qualitatively special views. We measured pose discrimination for a wide range of objects, finding large variations in performance depending on the object and the view angle, with front and back views yielding particularly good discrimination. Strikingly, a simple and biologically plausible computational model based on measuring the projected 3D optical flow between views of objects accurately predicted both successes and failures of discrimination performance. This provides a computational account of why certain views have a privileged status.

**Significance statement:** Some viewpoints of objects are qualitatively and perceptually special, making them easier to recognize and remember. We show that qualitatively special viewpoints of familiar and novel 3D objects can be predicted by an optical-flow model that measures how points on the surface shift in the image as viewpoint changes. This provides a quantitative account for why some viewpoints of objects are perceptually special.

## Introduction

When asked to imagine a familiar object, most people find themselves picturing the object from particular viewpoints that are especially informative or qualitatively distinct from other views (Palmer et al., 1981). Yet, despite decades of research on visual object perception, fundamental questions remain about why certain views of objects are special. There is a confusing array of terms and ideas related to different kinds of views, including ‘canonical’ (Blanz et al., 1996; Center et al., 2022; Cutzu & Edelman, 1994; Gomez et al., 2008; Woods et al., 2008), ‘accidental’ or ‘non-accidental’ (Koning & Lier, 2006; Niimi & Yokosawa, 2006; Poggio & Vetter, 1994), ‘generic’ (Freeman, 1994; Yuille et al., 2000, 2003), or ‘cardinal’ (or being aligned along a cardinal axis (Aldegheri et al., 2023; Appelle, 1972; Balas & Valente, 2012; Oomes & Dijkstra, 2002; Shiffrar & Shepard, 1991)). Here, we sought a computational framework for understanding why some views are privileged in object perception. We show that a simple computational model, based on optical flow (Stewart, Hartmann, et al., 2022) can accurately predict the costs and benefits of viewing both familiar and novel objects from certain perspectives. The model provides a straightforward quantitative account of how the visual system determines which views of objects are particularly important, based on regularities in object geometry, and the 2D visual information that is projected onto the retina.

Viewpoints play an important role in object perception, and can influence how we perceive, remember, and recognise an object. Here we will consider three specific, geometrically and qualitatively distinct viewpoints: (1) canonical viewpoints (Marr & Nishihara, 1978), which tend to be the oblique views where the most surface of the object is visible (Blanz et al., 1996; Bülthoff & Edelman, 1992); (2) end-on cardinal viewpoints, when an object is viewed so that the viewpoint with the smallest width to length ratio is aligned along the viewing axis (for example viewing a pig front-on); and (3) conversely the flat sides of objects, where the largest width to length ratio is aligned along the viewing axis (like the side of a pig) (Niimi & Yokosawa, 2008; Tarr & Kriegman, 2001). Viewing an object from one of these special viewpoints can benefit object recognition and recall (Center et al., 2022; Gomez et al., 2008; Palmer et al., 1981; Woods et al., 2008), and can result in longer inspection times of the object (Perrett & Harries, 1987). For example, the canonical viewpoints of an object might convey the most information about its overall identity and are the best views for object recognition (Marr & Nishihara, 1978), and conversely the flat viewpoints are the most perceptually stable, or provide the least amount of visual change if the object were to rotate a little (Niimi & Yokosawa, 2008; Tarr & Kriegman, 2001). People are also better at performing viewpoint discrimination tasks from some viewpoints. Previous work has attempted to quantitively define the transition between qualitatively distinct views (Tarr & Kriegman, 2001) as “visual events,” and found that when an object is rotated across one of these visual events, discrimination performance is higher. Such discrimination benefits were found to be particularly strong for the front and back of familiar objects, particularly when the objects were symmetrical, and/or had orientation-specific features such as linear contours (Niimi & Yokosawa, 2008).

We reasoned that we could use these geometric regularities in object viewpoint discrimination to provide a quantitative prediction of qualitatively distinct viewpoints, derived from the extent to which points on the object shift in the image when viewpoint (or equivalently, object pose) changes. To do this, we used a simple model inspired by optical flow computations, which was recently shown to capture the non-uniformities and geometric regularities in object viewpoint perception (Stewart, Hartmann, et al., 2022). Specifically, the model assumes that given a pair of views of an object, the visual system: (1) identifies corresponding points on the objects’ surface across the two poses, (2) estimates the vectors in 3D between these corresponding points; (3) projects the vectors into the image plane from the current perspective and (4) takes the average length of these vectors as a measure of the difference between the two poses. We find that this approach provides a quantitative account of the differences between views and thus a quantitative framework for understanding what makes certain object viewpoints qualitatively special.

## Results

### Experiment 1: viewpoint discrimination judgements

We reasoned that if the proposed optical flow model can be used to unify previous qualitative findings on cardinal viewpoints, it should first be able to predict discrimination benefits at cardinal (front and back) versus non-cardinal (oblique) viewpoints (Niimi & Yokosawa, 2008; Tarr & Kriegman, 2001). In two online experiments, we collected human discrimination judgements for cardinal and non-cardinal viewpoints for twenty-one photographs of real objects from the Amsterdam Library of Object Images (Geusebroek et al., 2005), and thirteen rendered mesh objects (see **Materials and Methods**). In an initial object priming block, participants were shown a video of each of the objects rotating for four seconds, and were asked to report the direction of rotation. This task aimed to prime participants to think about the 3D rotational nature of the objects in the subsequent task. Results from this block were only used for participant exclusion (one participant for this criterion), and were not analysed further. In the subsequent object discrimination block, two viewpoints of the same object were displayed either side of a central cross for 500ms: the base viewpoint was either a cardinal or non-cardinal viewpoint, and the rotated viewpoint was offset from the base view by seven possible rotation levels (0, ±5, ±10, ±15 degrees rotation around the vertical axis). Participants indicated whether the two viewpoints were the same or different by clicking an onscreen button (Figure 1A).

**Figure 1.**
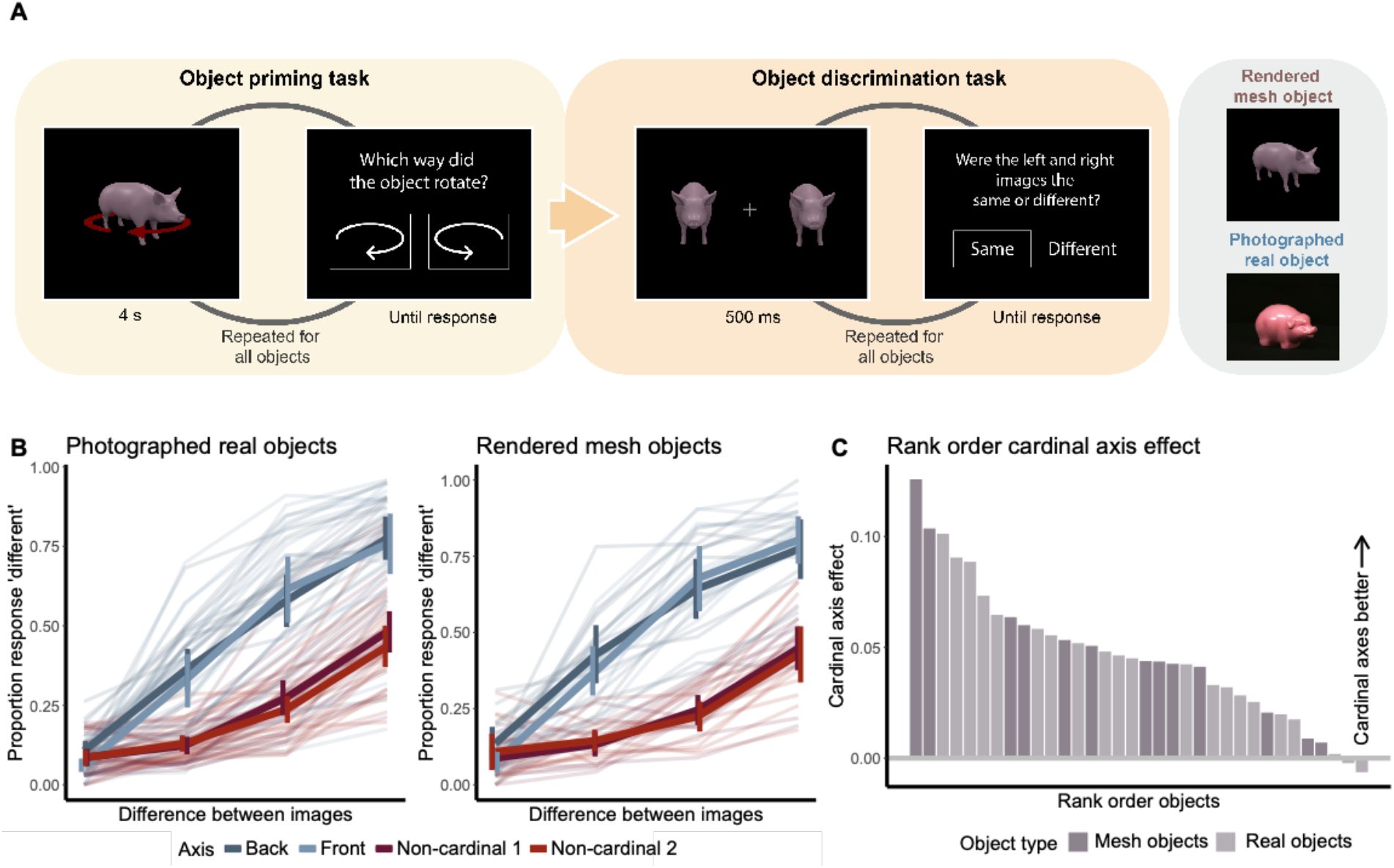
A) Participants first completed a priming task: each object rotated for 4 seconds; participants indicated rotation direction via button press. Participants then completed a pose discrimination task. Two viewpoints of an object were shown: the cardinal/non-cardinal viewpoint, and a viewpoint offset by 0, +- 5, 10, 15 degrees rotation. Participants had to indicate whether the two viewpoints were the same or different. Photographed real objects and rendered mesh objects were tested in two separate experiments. B) Human discrimination performance for photographed (left) and rendered (right) objects. Pale lines: individual participants; bold lines: mean; error bars indicate standard errors. C) The cardinal axis effect for each object, rank ordered. This was calculated as the slope between performance at 0- and 5-degrees difference. Real objects are depicted in light purple, mesh objects in dark purple.

To analyse the responses, for each object and rotation level, we calculated the proportion of responses where participants indicated the two views were the same (Figure 1B). Results showed a striking difference in performance between cardinal versus non-cardinal axes for both the photographs of real objects, and the rendered mesh objects. A generalized least squares (GLS) regression model (performance ∼ rotation_level (0, 5, 10, 15) * axis_type (cardinal, non-cardinal) * image_set (real, rendered)) demonstrated a significant effect of rotation level: F(1,536) = 734.76, p<0.0001; axis type: F(1,536) = 127.88, p<0.0001; and the interaction between rotation level and axis type: F(1,536) = 28.92, p<0.0001; but not image set: F(1,536) = 0.24, p = 0.63; and no other interactions were significant. This demonstrates that humans are better at discriminating objects rotated around cardinal versus non-cardinal axes for both real and rendered objects. These cardinal axes would therefore seem to be analogous to the “visual events” postulated by Tarr and Kriegman (Tarr & Kriegman, 2001).

The results also showed that there was some variability in performance between objects. To quantify the discrimination benefit for cardinal vs non-cardinal axes for a particular object, we calculated the “cardinal axis effect” as the difference in the slope between “response different” judgements for 0- and 5-degree rotation levels for cardinal vs non-cardinal axes, as this was the difference at which the most variability was observed between objects. The higher the slope for an object/axis, the more discriminable it is. Figure 2C shows that the cardinality effect was stronger for some objects than others, for both real and mesh objects, with 32/34 objects showing a perceptual discrimination benefit for cardinal versus non-cardinal viewpoints. This variability in the cardinal axis effect allowed us to use a model to investigate to what extent we could predict the cardinal axis effect for each object, and whether features of our model predictions might predict the magnitude of this effect.

**Figure 2.**
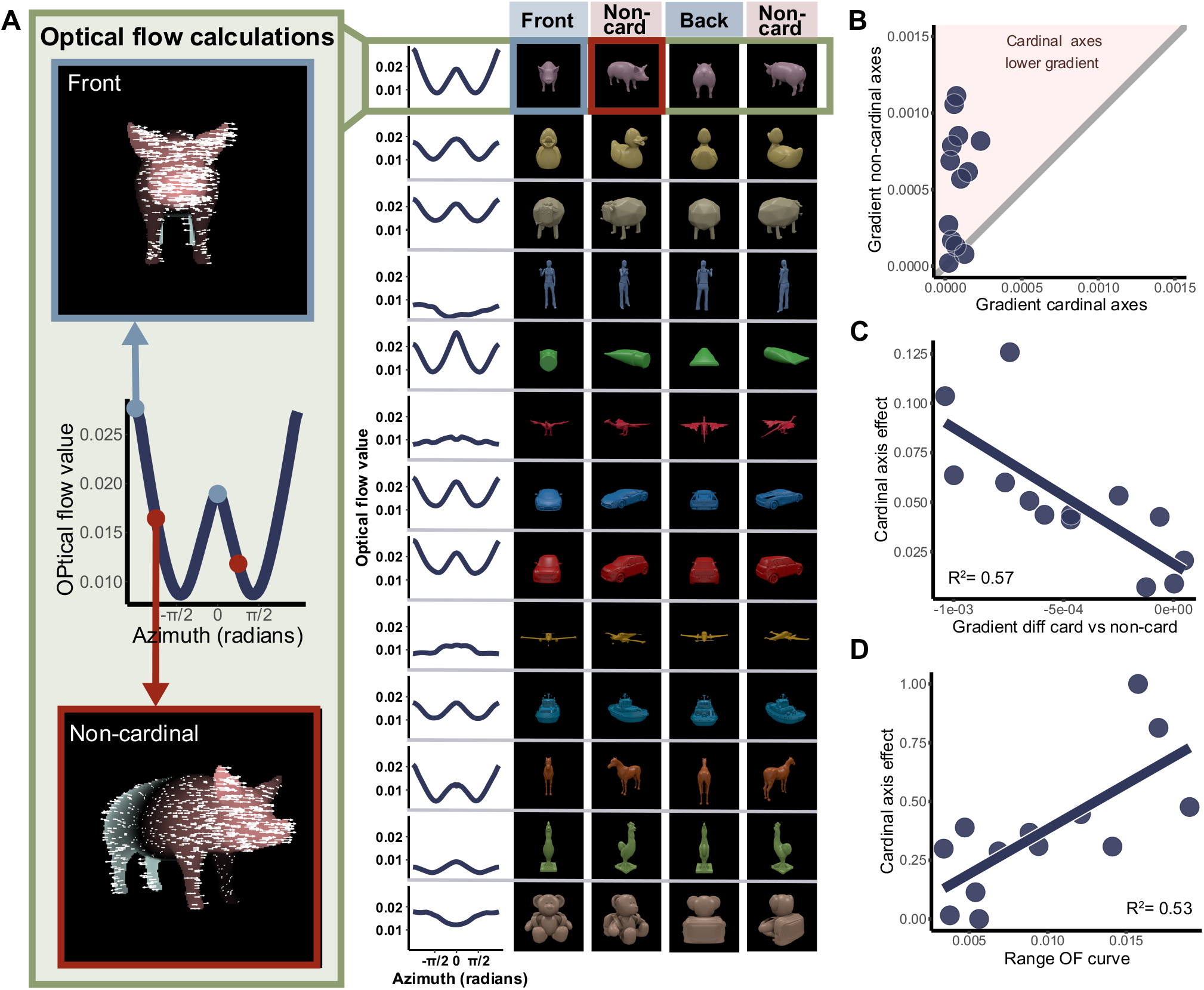
A) Optical flow model. For each view of each mesh object, the optical flow was calculated between that view and the next view, resulting in an optical flow curve (left, middle). Left top and bottom show example optical flow output for the Front and Non-cardinal axes of one object (pink indicates rightwards motion, green, leftwards). Right panel shows optical flow curves for each object, with front, back, and non-cardinal viewpoints of each object. B) Comparison of the gradient of cardinal versus non-cardinal axes on the optical flow curve. Dots represent individual objects. The pink shaded area represents objects where the optical flow gradient was lower for cardinal than non-cardinal axes. C) The cardinal axis effect (difference in slope between 0 and 5 degrees offset, as in Figure 1C) was predicted by the gradient difference between cardinal and non-cardinal axes on the optical flow curve. D) The cardinal axis was predicted by the range of the optical flow curve.

### Optical flow model predicts cardinal viewpoints

We used an optical flow model to compute viewpoint dissimilarity for every viewpoint around each object. In brief, the model measures how much points on objects shift in the image as the viewpoint changes. Specifically, given a pair of poses of an object, we compute the 3D vectors between corresponding surface points, and then estimate the mean length of these vectors when projected into the 2D image plane. This model has been shown to capture viewpoint-related variations in mental rotation (Stewart et al 2022), and we reasoned that it may be able to predict both cardinal and non-cardinal axes within objects, as well as the relative degree of the cardinal axis effect across objects. For each of the 72 rendered viewpoints, we calculated the ground-truth optical flow vectors produced as the object rotated towards the next viewpoint. The model prediction for this viewpoint was taken as the mean of the absolute length of these vectors (see Stewart et al for further details). We thus obtained an “optical flow curve” for each object (Figure 2). We calculated the gradient of this flow curve at the tested viewpoints, and the slope of the curve between the tested base viewpoint (front, back, non-cardinal) and each offset viewpoint. Different objects had different optical flow curve profiles, and in particular varied in the range of the curve. We therefore also calculated the range (max-min) of the curve for each object.

The gradient of the optical flow curve predicted cardinal versus non-cardinal axes, with cardinal axes having a lower gradient than non-cardinal axes (t(12) = -4.58, p=0.0006, Cohen’s D = 1.27 (large effect size), Figure 2AB). The optical flow model could also predict human discrimination performance in two ways. We first looked at whether the cardinal axis effect could be predicted by the gradient of the optical flow curve (Figure 2C). A simple linear regression model revealed that the model gradient accurately predicted the human discrimination performance (F(1,50) = 14.09, p = 0.00045). Second, we tested whether the cardinal axis effect could be predicted by the difference in gradient (Figure 2D), and by the magnitude of optical flow change across the entire optical flow curve (range of the curve). A linear regression model (cardinal_axis_effect ∼ gradient_difference + curve_range) again showed a significant effect of gradient difference (F(1,49) = 9.62, p = 0.0032) and curve range (F(1,49) = 56.2, p<0.0001). These results indicate that the gradient of the optical flow curve is predictive of whether a viewpoint is cardinal or not, providing for the first time a straightforward quantitative predictor for these qualitatively special views of familiar objects.

### Experiment 2: the front of novel objects

The objects tested in Experiment 1 were all easily recognizable objects, and factors such as familiarity and the geometric properties of the shapes themselves (e.g., symmetry, elongation), might have influenced performance. Thus, while model predictions correlated with the cardinal axis effect for symmetrical, elongated, real objects, the findings may not generalise to novel objects with less regular elongation and symmetry (Poggio & Vetter, 1994). We therefore created ten non-meaningful mesh objects with both regular and irregular optical flow curves, and in an online experiment verified that these objects were on average rated as non-familiar (see **Methods and Materials** for details of object-familiarity ratings). Objects were created to have varying levels of symmetry and elongation, and to have a more heterogenous pattern of optical flow predictions than the familiar objects in Experiment 1. We then conducted a separate online experiment, and asked a new sample of fifty online participants to rotate each of the ten novel objects, plus four of the familiar mesh objects from the previous experiment (pig, duck, small car, figure), so that the front of the object was facing toward the participant. For each object, we examined where the responded “front” angles lay on the optical flow prediction curve. In general, responses tended to cluster around the peaks and troughs of the optical flow curve (Figure 3), and on average the viewpoints that were indicated as being a “front” had lower gradients than the viewpoints that were never indicated as being the front (Wilcoxon test *z* = 2, p = 0.00037, *r* = 0.54; strong effect size). This demonstrates that, while there is naturally more variability in where the actual front of the object is considered to be, even for novel, non-symmetrical objects, the optical flow model was predictive of which viewpoints may be considered to be candidate “front” views of these objects.

**Figure 3.**
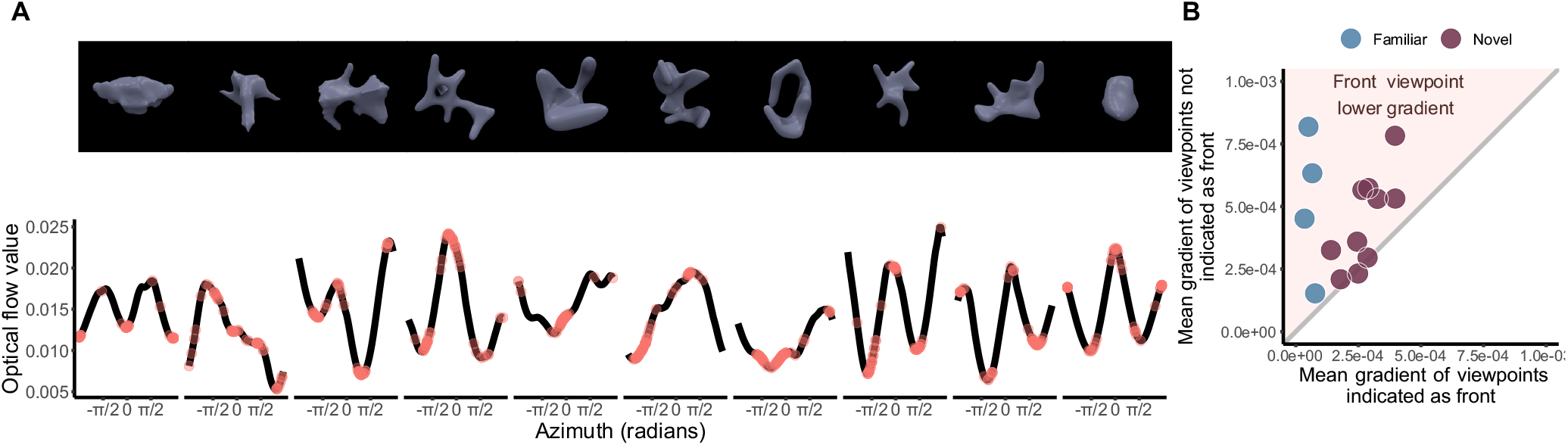
A) Tested novel objects (top) with the corresponding optical flow curve (bottom). Novel objects are shown from the viewpoint with the highest number of “front” responses. Individual points on the curve represent viewpoints that were marked as the “front” of the object. B) Scatter plot representing for each object the mean gradient across participants at viewpoints selected as “front” compared to viewpoints that were not selected as “front”. Novel objects are represented in purple, and the four familiar objects tested are shown in blue for comparison. The shaded pink area represents where the viewpoints selected as “front” have a lower gradient than non-front viewpoints.

## General discussion

Our results suggest that the optical flow model can explain variability in viewpoint discrimination and identify qualitatively distinct viewpoints. Participants were better at discriminating between two viewpoints separated by 5 degrees rotation, when one of those viewpoints was a so-called cardinal axis, compared to when it was an oblique view of the object. The magnitude and variability in this perceptual discrimination advantage could be predicted by the gradient of an optical-flow model that computes the magnitude of the 2D displacement vectors that would be produced if the object were to rotate from one viewpoint to the next. Remarkably, this model could also predict viewpoints that were more likely to be labelled as “front” for novel, unfamiliar objects. This model can therefore capture quantitative geometrical relationships between viewpoints and predict which viewpoints stand out as particularly significant for the observer. Our findings suggest the method works for familiar, unfamiliar, regular, and irregular objects. As Figure 4 shows, viewpoints that may be considered the most discriminable (front, back), most stable (sides), and “typical”, “generic” or “representative” (oblique; see Supplementary Materials) can be constrained using the model output curve (optical flow value/predicted perceived dissimilarity), and the gradient of this curve. Thus, a simple, quantitative account based on the projected spatial shifts of visible surface points provides a computational framework for object viewpoint perception.

**Figure 4.**
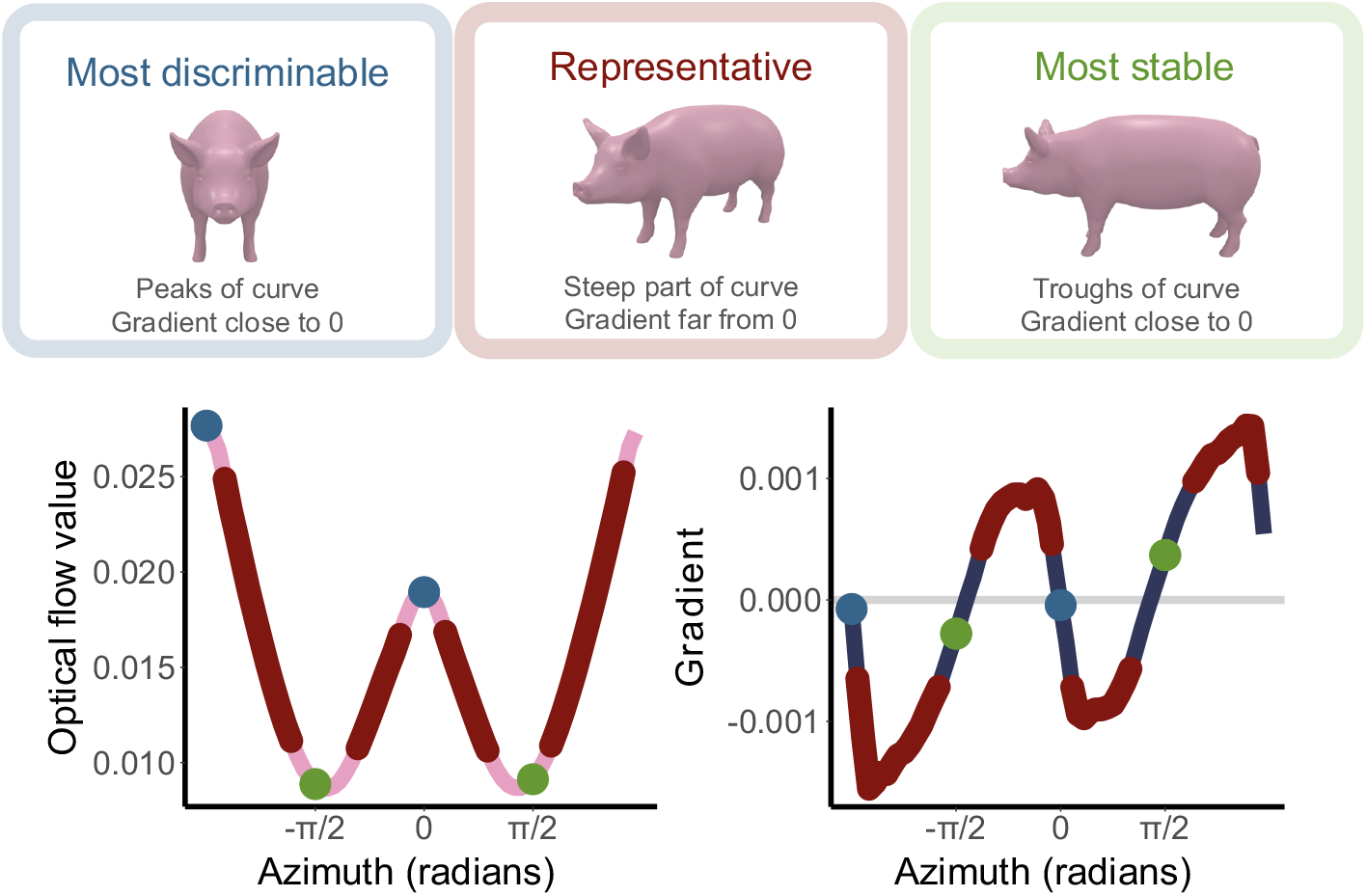
This model can quantitatively predict and constrain which viewpoints of a 3D object are qualitatively meaningful using the predicted dissimilarity curve, and the gradient of this curve.

It is likely that geometrical regularities across natural objects contribute to a form of statistical learning about object geometries. For example, in most quadrupedal animals, the front and back of the animal is narrower and more symmetrical than the side-on view. While such statistical learning about object categories and identity may account for familiarity effects in object recognition (Blanz et al., 1996), learning about the geometry of objects may also aid in identifying important viewpoints for unfamiliar objects. The results of Experiment 2 suggest that participants may extrapolate learned geometrical regularities of cardinal viewpoints experienced in the real world, and use these statistical regularities to determine the cardinal viewpoints of novel objects. As with many other object features (Kanade, 1981; Langer & Bülthoff, 2000; Mamassian & Landy, 2001; Sprote & Fleming, 2013; Wilder et al., 2011), the visual system seems to use knowledge of objects in the world to form priors about (Kersten et al., 2004), or estimate latent variables underlying (Fleming & Storrs, 2019), the proximal information about the geometry of distinctive object viewpoints.

These results also allow us to reflect on the nature of object viewpoint representations in relation to their true distal form as opposed to our 2D proximal experience of them. We recently suggested, based on the model also used in the current study, that the computations underlying the 3D mental rotation of objects rely on a ‘mental rendering’ of the imagined object, as if predicting its 2D proximal appearance (Stewart, Hartmann, et al., 2022). We suggest here that the comparison and categorisation of 3D object viewpoints might then rely on similar 3D to 2D computations. ^10^Interestingly, in this experiment, even though participants were primed on the 3D nature of the stimulus by being shown the object rotating in space, the model based on distances in 2D is still predictive of human performance, and reflects previous findings that a 2D representation may underpin 3D viewpoint discrimination (Stewart, Hartmann, et al., 2022). In Experiment 2, the selection of the “front” viewpoint compared to all other viewpoints arguably requires a 3D representation of the object as a whole, yet even in this case responses corresponded to geometric regularities predicted by the 2D model. As a result, the model also provides a route into understanding the origin of putative effects of the ‘perspectival’ appearance of objects (Morales et al., 2020; Storrs & Arnold, 2013; Thouless, 1931)— such as the perceived ‘elliptical nature’ of a coin seen slanted in depth—in the context of theories of vision that assume perceptual constancies (Burge, 2010; Cohen, 2014; Green, 2023; Walsh & Kulikowski, 1998). Specifically, we suggest that even when 3D object structure is estimated perfectly, comparisons between objects are made in terms of the estimated or predicted changes in the proximal stimulus involved in transforming one view to another. We speculate that representations that express changes in terms of proximal stimulus quantities are particularly useful in learning to see a 3D object in the absence of ground-truth data about its physical state. Specifically, we suggest that learning to accurately predict changes in the proximal stimulus teaches the visual system deep knowledge about the distal forms that produce such changes (Fleming & Storrs, 2019; Storrs & Fleming, 2021). Thus paradoxically, retaining a representation of ‘perspectival aspects’ of objects may be a key step in learning to see them as 3D in the first place. Our findings suggest that humans use proximal information about an object’s geometry to make judgements about an object’s pose, and more importantly, use this information to learn regularities about qualitatively special viewpoints.

## Materials and methods

### Participants

300 participants completed the study in total. Experiment 1 real objects: 50 participants (24 female; age range 19-59, mean age 29.9 (SD 1.3)), Experiment 1 rendered mesh objects (N = 50, female = 27, undisclosed sex = 1, mean age = 31.4 (SD 10.4)). Experiment 2 familiarity ratings 1 (N = 50, 18 female, mean age = 27.8 (SD 5.9)); familiarity ratings 2 (N = 50, 16 female, mean age = 27.9 (SD 8.5)). Experiment 2 front judgements: 50 participants (N = 50, 19 female, mean age = 30 (SD 8.7)). Experiment 2 representative judgements: 50 participants (N = 50, 28 female, mean age = 41.9 (SD 12.7)). Participants participated online, and were recruited through Prolific. Experiments were approved by the University of Marburg local ethics committee (approval number 2015-35k), and University of Giessen local ethics committee (approval number 2020-0033), and were conducted in accordance of the Declaration of Helsinki (1964).

### Stimuli

#### Experiment 1 – photographed real objects

Photographed real objects were taken from the Amsterdam Library of Object Images (ALOI) (Geusebroek et al., 2005). This library contains 1000 photographs of real-world images, photographed from 72 viewpoints separated by 5 degrees of horizontal rotation. In a previous study, we collected human judgements about which viewpoints of these objects corresponded to a number of axis labels (front, back, left, right, prototypical; see (Stewart, Ludwig, et al., 2022) for details on data collection). This data gave us a distribution of angular responses for each axis label for each object. For this study, we chose 21 objects that had clearly defined cardinal viewpoints: for each object and viewpoint label, we calculated the mean resultant length of participant responses, and used this to select objects where there was high agreement between participants for all cardinal viewpoints. The cardinal viewpoint for a specific label was then taken as the circular mode of the responses for that label. Non-cardinal viewpoints were taken as those that were midway between two cardinal viewpoints (i.e., front and left), but were not reported to be the prototypical or any other viewpoint.

#### Experiment 1 – rendered mesh objects

We repeated the experiment with mesh objects so that we could apply the mesh-based optical flow model. We selected 13 mesh objects that were as similar as possible to the original ALOI objects in both shape and semantic meaning. Meshes were either freely available online or selected from Evermotion (https://evermotion.org). These meshes were rendered in Blender (BlenderFoundation, 2018) from 72 viewing angles separated by 5 degrees horizontal rotation, using analogous lighting and camera distance as the ALOI objects (Geusebroek et al., 2005). For these objects, the front was defined by the researchers (and was analogous to the front of the photographed objects), and the same angular rotations relative to the front as in the photographed objects were used to define non-cardinal viewpoints. Objects were rendered in pretty colours chosen by the researcher.

#### Experiment 2 – unfamiliar objects

We created thirty semantically non-meaningful mesh objects using Mathematica and Blender (BlenderFoundation, 2018), and chose the ten most unfamiliar (see procedure for details) These objects were rendered from the same viewing angles, with the same lighting conditions as in Experiment 1. The four familiar objects from Experiment 1 were re-rendered in the same colour as the unfamiliar objects.

### Procedure

#### Experiment 1

Experiment 1 (photographed real objects) was pre-registered at as.predicted.org (#67124), and the experiment using the rendered mesh objects followed the same protocol. The experiments were conducted online and programmed with custom-written software using JsPsych (deLeeuw, 2015). Each participant completed both an object priming and a perceptual discrimination task. The object priming task aimed to familiarise participants with the 3D nature of the objects, and to prime them to think about the objects rotating through viewpoints. Participants viewed a video of each object rotating through 360 degrees for a duration of four seconds, either clockwise or counter-clockwise. Participants had to indicate the direction of rotation via button-press. Participant responses to the priming task were only used for excluding participants and were not analysed further. In the perceptual discrimination task, participants were presented with two views of an object on either side of a central fixation cross, for 500ms. They then indicated via button press whether the views were the same or different. These views were defined by four conditions, tested in two separate experiments. Experiments 1 and 2 (part 1): front ± 0, 5, 10, 15 degrees; non-cardinal angle 1 ± 0, 5, 10, 15 degrees. Experiments 1 and 2 (part 2): back ± 0, 5, 10, 15 degrees; non-cardinal angle 2 ± 0, 5, 10, 15 degrees. Every participant saw all objects at both cardinal and non-cardinal viewpoints, with randomized viewpoint offset difference levels.

### Experiment 2

#### Familiarity ratings

To choose Participants saw a video of each object rotating, and indicated how familiar they found the object to be, using a 6-point slider ranging from “very unfamiliar” to “very familiar”. The video looped to give the appearance of a continuously rotating object until a response was given. Four of the objects from experiment 2 were included as catch and comparison trials (pig, car, figure, duck). Mean familiarity ratings were calculated, and objects were rank ordered by familiarity. We chose 10 objects for the cardinal axis rating task, accounting for low familiarity ratings, varying levels of symmetry, and varying optical flow curve profiles. To confirm that these chosen novel objects were perceived as being unfamiliar, we ran a second online experiment with the same procedure, where participants only saw these ten unfamiliar objects and the four familiar objects (see Supplementary Materials). *Cardinal axis ratings*. In an online experiment, participants were instructed to rotate each object using the left and right arrows on the keyboard until the object was facing toward the “front”.

### Exclusions

In Experiment 1, trials were excluded if the reaction time was less than 300ms or more than 5000ms or if performance was less than 75% correct on either the training or main task. In Experiment 2, for the familiarity ratings, trials were excluded if the reaction time was less than 300ms or more than 5000ms, or if they rated any of the real objects (pig, figure, car) as having a familiarity rating of less than three out of six. For the front ratings, trials were excluded based on the same reaction time criteria, and additionally if the “front” response of familiar objects was not within 45 degrees of the veridical front. 1.2% of trials were excluded for Experiment 1 (photographed real objects) and 1.19% for the rendered mesh objects. For Experiment 2 (familiarity ratings) no trials were excluded; for Experiment 2 (front ratings), 3 participants were excluded, and no trials were excluded from the remaining participants.

### Analyses

#### Model

We used a ground-truth optical flow computation (Stewart et al, 2022), which has previously been found to predict human performance for viewpoint dissimilarity judgements for block-sequence 3D rendered objects. The model predicted the amount of 2D optical flow information that would be produced as the object rotated by 5 degrees from one viewpoint to the next, by calculating the mean of the absolute horizontal and vertical displacements of every visible vertex in the underlying object mesh. Displacements for vertices visible from one viewpoint but not the rotated viewpoint (unmatched points) are not included in model calculations.

#### Cardinal axis effect

To calculate the cardinal axis effect, we only took the slope of performance difference between 0- and 5-degree displacements for the following reasons: 1) this is the most difficult discrimination level we tested; 2) larger discrimination levels (especially 15-degree become trivial); 3) there’s not necessarily a linear increase in performance between 0,5-,10-, and 15-degree displacements. For some objects there is a very small performance difference between 0- and 5-degree displacements; and for others there is a larger difference in performance. Such between-object differences in sensitivity to small displacements are lost if larger displacements are included in the cardinal axis effect calculations.

#### Statistical analyses

All statistical analyses were conducted in R. Linear regression models were conducted using base R. Generalized least squares (GLS) models were conducted using the package nlme (Pinheiro et al., 2020), and were implemented where a linear model would otherwise have a heterogonous variance across different levels of a factor. Model comparisons were used to determine the best-fitting variance structure (different variance allowed for different factor levels).

## Acknowledgements

This project was supported by the Deutsche Forschungsgemeinschaft, through project numbers 460533638 (EEMS) and 222641018–SFB/TRR-135 TP C1 (RWF) and TP B2 (AS), by the Research Cluster “The Adaptive Mind”, funded by the Hessian Ministry for Higher Education, Research, Science and the Arts and by the European Research Council through projects ERC-2022-AdG “STUFF” (project number 101098225 to RWF) and ERC-2020-CoG “SENCES” (project number 101001250 to AS). Data and stimuli will be made available upon publication at http://doi.org/10.5281/zenodo.8398391.

## References

Aldegheri, G., Gayet, S., & Peelen, M. V. (2023). Scene context automatically drives predictions of object transformations. Cognition, 238, 105521. 10.1016/j.cognition.2023.105521

Appelle, S. (1972). Perception and discrimination as a function of stimulus orientation:The “oblique effect” in man and animals. Psychological Bulletin, 78(4), 266–278. 10.1037/h0033117

Balas, B., & Valente, N. (2012). View-adaptation reveals coding of face pose along image, not object, axes. Vision Research, 67, 22–27. 10.1016/j.visres.2012.07.002

Blanz, V., Tarr, M. J., & Bülthoff, H. H. (1996). What Object Attributes Determine Canonical Views? Perception, 28(5), 575–599. 10.1068/p2897

BlenderFoundation. (2018). Blender-a 3D modelling and rendering package (http://www.blender.org).

Bülthoff, H. H., & Edelman, S. (1992). Psychophysical support for a two-dimensional view interpolation theory of object recognition. Proceedings of the National Academy of Sciences, 89(1), 60–64. 10.1073/pnas.89.1.60

Burge, T. (2010). Origins of Objectivity. 137–153. 10.1093/acprof:oso/9780199581405.003.0005

Center, E. G., Gephart, A. M., Yang, P.-L., & Beck, D. M. (2022). Typical viewpoints of objects are better detected than atypical ones. Journal of Vision, 22(12), 1. 10.1167/jov.22.12.1

Cohen, J. (2014). Perceptual Constancy. In M. Matthen (Ed.), The Oxford Handbook of Philosophy of Perception (pp. 621–639). 10.1093/oxfordhb/9780199600472.001.0001

Cutzu, F., & Edelman, S. (1994). Canonical views in object representation and recognition. Vision Research, 34(22), 3037–3056. 10.1016/0042-6989(94)90277-1

deLeeuw, J. R. (2015). jsPsych:A JavaScript library for creating behavioral experiments in a Web browser. Behavior Research Methods, 47(1), 1–12. 10.3758/s13428-014-0458-y

Fleming, R. W., & Storrs, K. R. (2019). Learning to see stuff. Current Opinion in Behavioral Sciences, 30, 100–108. 10.1016/j.cobeha.2019.07.004

Freeman, W. T. (1994). The generic viewpoint assumption in a framework for visual perception. Nature, 368(6471), 542–545. 10.1038/368542a0

Geusebroek, J.-M., Burghouts, G. J., & Smeulders, A. W. M. (2005). The Amsterdam Library of Object Images. International Journal of Computer Vision, 61(1), 103–112. 10.1023/b:visi.0000042993.50813.60

Gomez, P., Shutter, J., & Rouder, J. N. (2008). Memory for objects in canonical and noncanonical viewpoints. Psychonomic Bulletin & Review, 15(5), 940–944. 10.3758/pbr.15.5.940

Green, E. J. (2023). Perceptual constancy and perceptual representation. Analytic Philosophy. 10.1111/phib.12293

Kanade, T. (1981). Recovery of the three-dimensional shape of an object from a single view. Artificial Intelligence, 17(1–3), 409–460. 10.1016/0004-3702(81)90031-x

Kersten, D., Mamassian, P., & Yuille, A. (2004). Object Perception as Bayesian Inference. Annual Review of Psychology, 55(1), 271–304. 10.1146/annurev.psych.55.090902.142005

Koning, A., & Lier, R. van. (2006). No symmetry advantage when object matching involves accidental viewpoints. Psychological Research, 70(1), 52–58. 10.1007/s00426-004-0191-8

Langer, M. S., & Bülthoff, H. H. (2000). A Prior for Global Convexity in Local Shape-from-Shading. Perception, 30(4), 403–410. 10.1068/p3178

Mamassian, P., & Landy, M. S. (2001). Interaction of visual prior constraints. Vision Research, 41(20), 2653–2668. 10.1016/s0042-6989(01)00147-x

Marr, D., & Nishihara, H. K. (1978). Representation and recognition of the spatial organization of three-dimensional shapes. Proceedings of the Royal Society of London. Series B. Biological Sciences, 200(1140), 269–294. 10.1098/rspb.1978.0020

Morales, J., Bax, A., & Firestone, C. (2020). Sustained representation of perspectival shape. Proceedings of the National Academy of Sciences, 202000715. 10.1073/pnas.2000715117

Niimi, R., & Yokosawa, K. (2006). Viewpoint Dependence in the Recognition of Non-Elongated Familiar Objects:Testing the Effects of Symmetry, Front-Back Axis, and Familiarity. Perception, 38(4), 533–551. 10.1068/p6161

Niimi, R., & Yokosawa, K. (2008). Determining the orientation of depth-rotated familiar objects. Psychonomic Bulletin & Review, 15(1), 208–214. 10.3758/pbr.15.1.208

Oomes, A. H. J., & Dijkstra, T. M. H. (2002). Object pose:Perceiving 3-D shape as sticks and slabs. Perception & Psychophysics, 64(4), 507–520. 10.3758/bf03194722

Palmer, S., Rosch, E., & chase, P. (1981). Canonical Perspective and the Perception of Objects. In J. Long & A. Baddeley (Eds.), Attention and Performance IX (pp. 135–151). Lawrence Erlbaum Associates.

Perrett, D. I., & Harries, M. H. (1987). Characteristic Views and the Visual Inspection of Simple Faceted and Smooth Objects:‘Tetrahedra and Potatoes.’ Perception, 17(6), 703–720. 10.1068/p170703

Pinheiro, J., Bates, D., DebRoy, S., Sarkar, D., & Team, R. C. (2020). nlme:Linear and Nonlinear Mixed Effects Models. https://CRAN.R-project.org/package=nlme

Poggio, T., & Vetter, T. (1994). Symmetric 3D objects are an easy case for 2D object recognition. Spatial Vision, 8(4), 443–453. 10.1163/156856894x00107

Shiffrar, M. M., & Shepard, R. N. (1991). Comparison of Cube Rotations Around Axes Inclined Relative to the Environment or to the Cube. Journal of Experimental Psychology:Human Perception and Performance, 17(1), 44–54. 10.1037/0096-1523.17.1.44

Sprote, P., & Fleming, R. W. (2013). Concavities, negative parts, and the perception that shapes are complete. Journal of Vision, 13(14), 3–3. 10.1167/13.14.3

Stewart, E. E. M., Hartmann, F. T., Morgenstern, Y., Storrs, K. R., Maiello, G., & Fleming, R. W. (2022). Mental object rotation based on two-dimensional visual representations. Current Biology, 32(21), R1224–R1225. 10.1016/j.cub.2022.09.036

Stewart, E. E. M., Ludwig, C. J. H., & Schütz, A. C. (2022). Humans represent the precision and utility of information acquired across fixations. Scientific Reports, 12(1), 2411. 10.1038/s41598-022-06357-7

Storrs, K. R., & Arnold, D. H. (2013). Shape AfteREFfects REFlect Shape Constancy Operations:Appearance Matters. Journal of Experimental Psychology:Human Perception and Performance, 39(3), 616–622. 10.1037/a0032240

Storrs, K. R., & Fleming, R. W. (2021). Learning About the World by Learning About Images. Current Directions in Psychological Science, 30(2), 120–128. 10.1177/0963721421990334

Tarr, M. J., & Kriegman, D. J. (2001). What defines a view? Vision Research, 41(15), 1981–2004. 10.1016/s0042-6989(01)00024-4

Thouless, R. H. (1931). PHENOMENAL REGRESSION TO THE ‘REAL’ OBJECT. II. British Journal of Psychology. General Section, 22(1), 1–30. 10.1111/j.2044-8295.1931.tb00609.x

Walsh, V., & Kulikowski, J. (1998). Perceptual Constancy:Why Things Look as They Do. Cambridge University Press. https://books.google.de/books?id=LAtJHcBRAkEC

Wilder, J., Feldman, J., & Singh, M. (2011). Superordinate shape classification using natural shape statistics. Cognition, 119(3), 325–340. 10.1016/j.cognition.2011.01.009

Woods, A. T., Moore, A., & Newell, F. N. (2008). Canonical Views in Haptic Object Perception. Perception, 37(12), 1867–1878. 10.1068/p6038

Yuille, A. L., Coughlan, J. M., & Konishi, S. (2000). The generic viewpoint constraint resolves the generalized bas relief ambiguity. Proc. Conf. Information Sciences and Systems.

Yuille, A. L., Coughlan, J. M., & Konishi, S. (2003). The generic viewpoint assumption and planar bias. IEEE Transactions on Pattern Analysis and Machine Intelligence, 25(6), 775–778. 10.1109/tpami.2003.1201826

