## Supplementary materials for "A simple optical flow model explains why certain object viewpoints are special"

**Supplementary methods.** *Representative viewpoints of novel objects.* We conducted an additional online experiment similar to Experiment 2, to test whether participant perception of the most representative view of an object corresponded to the higher gradient areas on the optical flow curve. We used the same objects as in Experiment 2, and following the methodology from Blanz et al. [1], participants (N = 50 after exclusions, 4 excluded for not performing any rotation; age = 41.9 (12.7), female = 28, 2.7% remaining trials excluded for the same criteria) were given the instructions to rotate the object to show the best impression of the object. Specifically: *"Imagine that you are making a brochure and are trying to give your customers the best possible impression of the object shown on the screen. Which view of the object would you choose?"*

**Supplementary results.** *Familiarity ratings for novel objects***.** To confirm that the objects used in the novel objects task were perceived as being unfamiliar, we rank-ordered familiarity ratings for the ten novel and four familiar objects used in these experiments (Figure S1). All novel objects were rated as being unfamiliar.

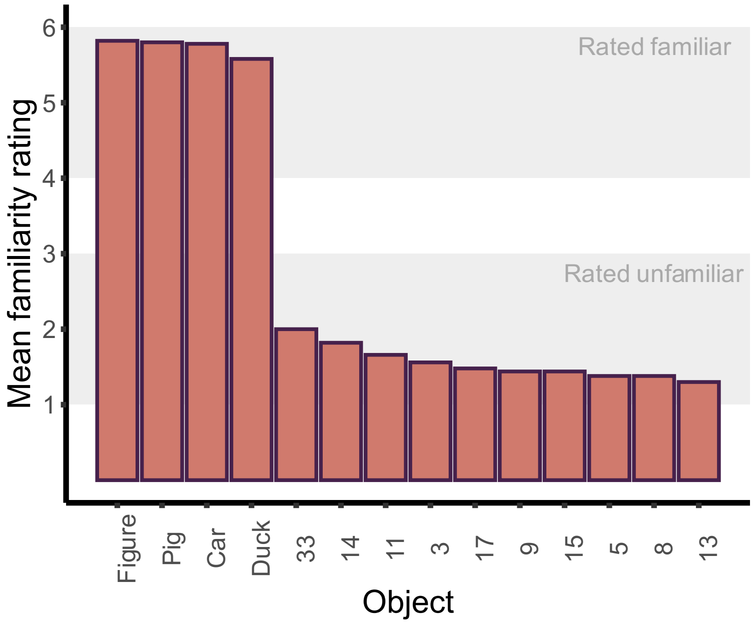


**Figure S1**. Rank ordered familiarity ratings for the ten novel and four familiar objects used in the novel object experiments.

*Representative views of novel object.* The viewpoints considered to be representative are labelled in blue in Figure S2, and it is evident that these viewpoints are shifted toward the higher-gradient slopes of the optical flow curve, compared to the viewpoints labelled as "front" (see Figure 4 in the main manuscript), and to more oblique viewpoints (demonstrated especially in the last four familiar objects). We compared the gradient at "front" versus "representative" viewpoints. For both familiar and unfamiliar objects, the front had a significantly lower gradient than representative viewpoints: Mann-Whitney U test, Familiar: W = 1990, $p < 0.0001$; Unfamiliar: W = 94079, $p < 0.0001$). As can be seen in Figure S3, the representative labelled viewpoints do not show as much clustering as the front labelled viewpoints, especially for unfamiliar objects. This is to be expected as, by definition, generic views of objects are not singular, so many different viewpoints may be considered representative. These results also reflect previous work that has shown less agreement in canonical viewpoints for unfamiliar compared to familiar objects [1]. Nevertheless, the results point to a tendency for the most representative view to be the oblique views that captures information about both the front and side of objects.

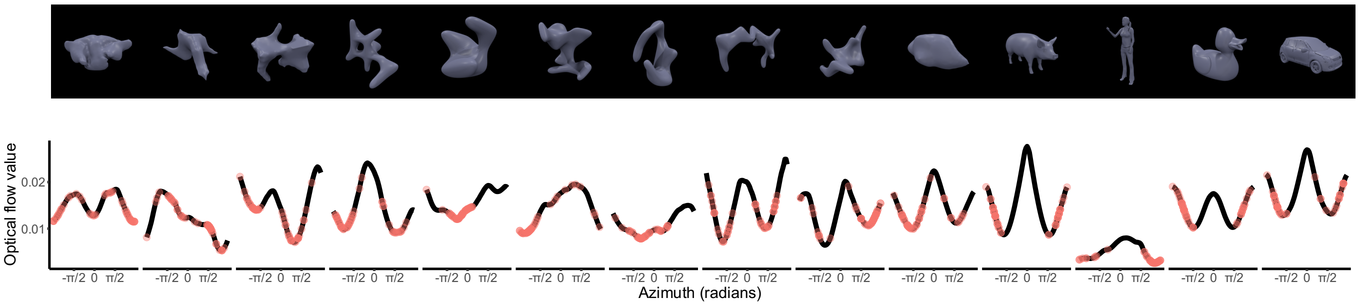


**Figure S2**. Tested objects (top) with the corresponding optical flow curve (bottom). Objects are shown from the viewpoint with the highest number of "representative" responses. Individual points on the curve represent viewpoints that were marked as "representative" of the object.

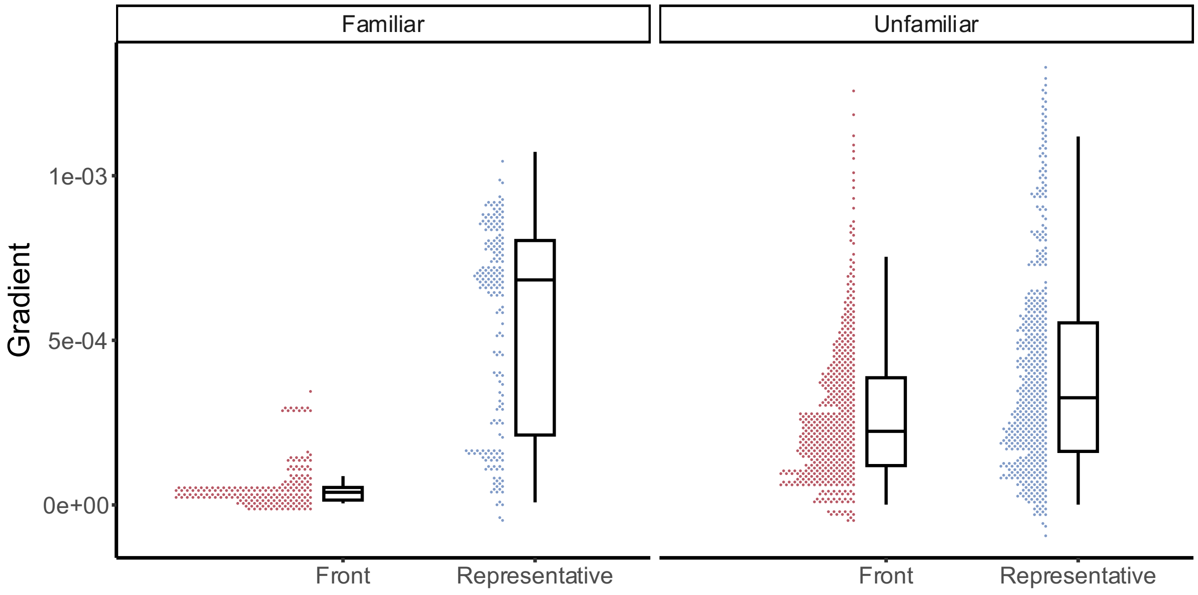


**Figure S3**. Median and IQR of viewpoints labelled as "front" and "representative" for both familiar and unfamiliar objects. Dot clouds visualise the distribution of responses.

**Supplementary references**


1. Blanz, V., Tarr, M. J. & Bülthoff, H. H. What Object Attributes Determine Canonical Views? *Perception* **28**, 575–599 (1996).
